# Esrrγa regulates nephron development and ciliogenesis by controlling prostaglandin synthesis and cooperation with Ppargc1a

**DOI:** 10.1101/2021.11.17.468871

**Authors:** Hannah M. Wesselman, Ana L. Flores-Mireles, Rebecca A. Wingert

## Abstract

Cilia are essential for the ontogeny and function of many tissues, including the kidney. In mammals, Esrrγ has been previously established as a significant determinant of renal health, with decreased expression linked to age related dysfunction, cyst formation, and kidney disease. Here, we report that the Esrrγ vertebrate ortholog *estrogen related receptor gamma a* (*esrrγa*) is essential for proper cell fate choice within kidney functional units (nephrons) as well as ciliogenesis. Deficiency of *esrrγa* resulted in nephrons with alterations in proximodistal segmentation and a decreased multiciliated epithelial cell populace. Surprisingly, *esrrγa* deficiency disrupted renal ciliogenesis and caused a similar abrogation within the developing node and otic vesicle—all defects that occurred independently of changes in cell polarity or basal body organization. These phenotypes were consistent with interruptions in prostaglandin signaling, and we found that ciliogenesis was rescued in *esrrγa* deficient embryos with exogenous PGE_2_ or through overexpression of the cyclooxygenase enzyme Ptgs1. Through genetic interaction studies, we found that peroxisome *proliferator-activated receptor gamma, coactivator 1 alpha* (*ppargc1a*), which acts upstream of Ptgs1-mediated prostaglandin synthesis, has a synergistic relationship with *esrrγa* in the ciliogenic pathway. These data position *esrrγa* as a novel link between ciliogenesis and nephrogenesis through regulation of prostaglandin signaling and cooperation with *ppargc1a*, and highlight *esrrγa* as a potential new target for future ciliopathic treatments.

## Introduction

Cilia are specialized surface organelles that are present on nearly every vertebrate cell where they serve critical functions including mechano- and chemo- sensing. In the case of motile cilia, primary cilia or multiciliated cells (MCCs) can facilitate fluid propulsion. During development, cilia are essential for the establishment and maintenance of planar cell polarity and organization of essential signaling molecules. For example, aberrant ciliogenesis results in disease states of several tissues, including the kidney, liver, pancreas, retina, and central nervous system (Pazour et al., 2019). Cilia defects have been linked to kidney disorders like polycystic kidney disease, Bardet-Biedl syndrome, Joubert syndrome, and many others (McConnachie et al., 2020). Etiologies for these conditions are varied, but many have been linked to mutations in a variety of specialized ciliary proteins. Production and maintenance of healthy cilia also require proper control of the ciliogenic transcriptional program. Such regulators include the RFX family of transcription factors, which are essential for both primary and motile cilia formation, and interactions with Foxj1 that can further regulate the development of motile cilia (Thomas et al., 2010). Furthermore, the hepatocyte nuclear factor 1B (HNF1B) regulates several ciliary genes, thereby contributing to kidney development and/or disease progression (Clissold et al., 2015, Gresh et al., 2004, Heisberger et al., 2005, Naylor et al., 2014, Sander et al., 2019, Thomas et al., 2010).

Recent studies have identified additional genetic networks that influence cilia formation. Among these, *peroxisome proliferator-activated receptor (PPAR) gamma, coactivator 1 alpha* (*ppargc1a*, known as PGC1*α* in mammals) regulates the ciliogenic program, in part, through control of prostanoid production (Chambers et al., 2018, Chambers et al., 2020). *ppargc1a* induces the biosynthesis of prostaglandin E_2_ (PGE_2_) by promoting the expression of *prostaglandin-endoperoxide synthase 1* (*ptgs1,* also known as *cyclooxygenase 1* (*cox1*)) in both the adult mammalian kidney and zebrafish embryonic kidney (Chambers et al., 2020, Tran et al., 2016). In turn, PGE_2_ is required for proper ciliary outgrowth by modulating intraflagellar transport (Jin et al., 2014). Interestingly, *ppargc1a* expression and PGE_2_ production are required for nephrogenesis, where they are involved in the pattern formation of nephron tubule segments and mitigate the fate choice between MCC and transporter cell lineages within these segments (Chambers et al., 2018, Chambers et al., 2020, Marra et al., 2019a, Poureetezadi et al., 2016).

Interestingly, ESRRγ, an orphan nuclear receptor, has been found to interact with both HNF1B and PGC1*α* in multiple contexts. ESRRγ and HNF1B cooperate in the regulation of mitochondrial function and proximal kidney cell development (Sander et al., 2019, Zhao et al., 2018). Similarly, ESRRγ and PGC1*α* bind common hormone response elements in kidney cells, and work synergistically in mitochondrial biogenesis (Liu et al., 2005, Fan et al., 2018, Wang et al., 2008). Phenotypes observed in *Esrrγ* knockout models further support its role in the regulation of energy production, as tissues with high energy demand, including the heart and kidney, are dysregulated. Specifically, *Esrrγ* knockout mice die soon after birth due to cardiac defects, and the renal tissue of these mice exhibits decreased ureteric branching (Alaynick et al. 2007, Berry et al., 2011). Furthermore, the kidney specific Esrrγ murine knockout results in kidney cysts and renal dysfunction (Zhao et al., 2018). Collectively, these phenotypes suggest that Esrrγ may play multiple roles in kidney and cilia development, yet neither analysis of nephron composition nor cilia formation has been reported to date.

Here, we report that *esrrγa* is necessary for proper nephron segmentation and ciliogenesis. In the zebrafish embryonic kidney, or pronephros, we found that deficiency of *esrrγa* resulted in segment patterning defects that included a decreased number of MCCs. Cilia projecting from MCCs and primary epithelial cells were also significantly shortened in both renal and non-renal populations within *esrrγa* deficient animals. These characteristics were strikingly reminiscent of insufficient prostaglandin signaling during development, and indeed we found that *esrrγa* ciliopathic phenotypes were rescued by supplementation of PGE_2_ or transcripts encoding the biosynthesis enzyme Ptgs1. Additionally, we discovered a synergistic genetic interaction between *esrrγa* and *ppargc1a* that is essential for renal cila formation. These findings provide fundamental new insights about the regulatory networks that direct ciliogenesis and renal development.

## Results

### *esrrγa* is expressed in renal progenitors

Prior research has demonstrated that *Esrrγ* is expressed in the mouse and human kidney, with particularly high expression profiles in the loop of Henle (RID: N-GK5G, 2-5CE6, 2-5CEA, 16- 5WSW) (Harding et al., 2011, Lindstrom et al., 2020, McMahon et al., 2008). Due to evolutionary whole genome duplication events, zebrafish have two homologs for Esrrγ—*esrrγa,* and *esrrγb* (Thome et al., 2014). Of these, only *esrrγa* is specifically expressed in a pattern consistent with its localization to renal progenitors and is later the distal nephron region, while *esrrγb* is not spatially restricted through early developmental stages (Bertrand et al., 2007, Thisse and Thisse, 2008).

To further assess the expression of *esrrγa* during pronephros ontogeny, we performed whole mount *in situ* hybridization (WISH) in wild-type (WT) embryos. Since renal progenitors are patterned into distinct segments by the 28 ss (**Figure 1A**), we assessed *esrrγa* expression between the 5 and 28 somite stage (ss). Transcripts were detected in the bilateral stripes of renal progenitors at the 8 ss, and continued to be expressed in nephron distal tubule segments at the 28 ss (**Figure 1B, Supplement 1D**). Fluorescent *in situ* hybridization at the 20 ss revealed that *esrrγa* transcripts colocalize with the essential kidney transcription factor *pax2a*, as well as nephron marker *cdh17* at the 28 ss (**Figure 1B**). Given this expression pattern throughout nephron development, we hypothesized that *esrrγa* may have roles in segment patterning and/or cellular differentiation.

**Figure 1.**
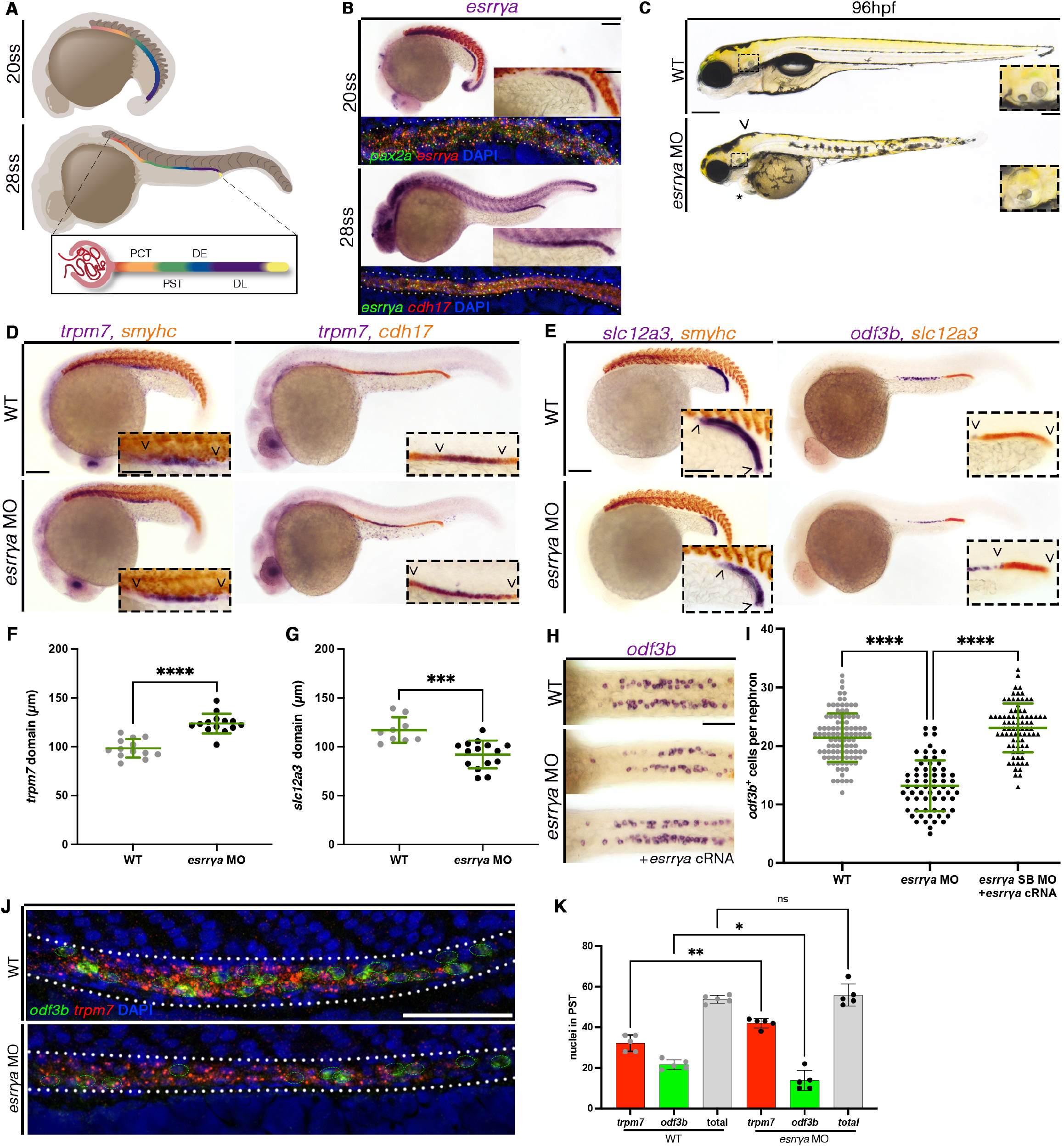
*Esrrγa* deficient animals exhibit altered morphology and nephron composition. (A) Schematic of nephrogenesis in a zebrafish from the early 20 ss to the fully patterned 28 ss. Nephron is comprised of the proximal convoluted tubule (PCT), proximal straight tubule (PST), distal early (DE) and distal late (DL) segments. (B) *esrrγa* is expressed in the kidney progenitors at the 20 ss (top) colocalizes with early marker (*pax2a*). Somites are marked with *smyhc* stain in red at the 20 ss. At the 28 ss (bottom) *esrrγa* colocalizes with the nephron marker (*cdh17*). Scale bar = 100µm for lower magnification image, scale bar = 50µm for both higher magnification images(C) WT sibling (top) and e*srrγa* morphant (bottom) zebrafish with pericardial edema (asterisk), hydrocephaly (arrow head), and fused otoliths (dashed box outline, inset). Scale bar = 100µm, inset = 50µm. (D) 20 ss (left) and 28 ss (right) WT (top) and *esrrγa* MO (bottom) animals stained via WISH for the PST marker (*trpm7*) with either somite marker (*smyhc*) or nephron marker (*cdh17*). Scale bar = 100µm for lower magnification, scale bar = 50µm for higher magnification. (E) 20 ss (left) and 28 ss (right) WT (top) and *esrrγa* MO (bottom) animals stained via WISH for the DL marker (*slc12a3*) with either somite marker (*smyhc*) or MCC marker (*odf3b*). Scale bar = 100µm for lower magnification, scale bar = 50µm for higher magnification. (F) PST domain length at 28 ss in micrometers. (G) DL domain length at 28 ss in micrometers. (H) WT (top) and *esrrγa* MO injected (middle and bottom), and *esrrγa* cRNA co-injected (bottom) zebrafish at 24 hpf stained with WISH for MCCs (*odf3b*). Scale bar = 50µm. (I) Absolute number of MCCs per nephron of zebrafish at 24 hpf. (J) Fluorescent in situ hybridization for MCCs (*odf3b*, green), and PST cells (*trpm7*, red), in WT and *esrrγa* MO injected zebrafish at 24 hpf. MCCs are circled in green dotted line. Scale bar = 50µm. (K) Absolute cell number of either *odf3b* or *trpm7* expressing cells in WT and *esrrγa* MO injected animals at 24 hpf. Data presented on graphs are represented as mean ± SD; ***p < 0.001 and ****p < 0.0001 (t test or ANOVA).

### *esrrγa* is required for nephron segmentation

In the developing mouse kidney, Esrrγ knockout disrupts branching morphogenesis, renal papilla formation, and causes perinatal lethality; additionally, kidney specific Esrrγ deficiency results in renal cyst formation (Berry et al., 2011, Zhao et al., 2018). To interrogate the function of *esrrγa* during zebrafish pronephros development, we performed loss of function studies using two previously published morpholinos (MOs) (Tohme et al., 2014). Of these, one morpholino was designed to block protein translation (*esrrγa* ATG MO), and the other was designed to interfere with splicing by blocking the exon 1 splice donor site (*esrrγa* SB MO) (Tohme et al., 2014) (**Supplement 1A**). The *esrrγa* SB MO caused improper splicing between exon 1 and 2 whereby a portion of exon 1 was excised in the process and produced a transcript that encoded a premature stop codon (**Supplement 1B-C**). Compared to WT embryos, *esrrγa* morphants displayed hydrocephaly, pericardial edema, and otolith malformations at 96 hours post fertilization (hpf) (**Figure 1C, Supplement 3A**). This combination of morphological phenotypes suggested that nephron segment development and/or ciliary function were compromised. Therefore, we next assessed the effect of *esrrγa* deficiency on each of the nephron segments and distinct ciliated cell populations within the kidney.

We found that upon knockdown of *esrrγa*, the proximal straight tubule (PST, marked by *trpm7*) expanded, while the distal late segment (DL, marked by *slc12a3*) decreased in length at both the 20 ss and 28 ss (**Figure 1D, 1E**). The proximal convoluted tubule (PCT, marked by *slc20a1a*), the distal early segment (DE, marked by *slc12a1*), and the overall length of the nephron tubule (marked by *cdh17*) remained unchanged (**Supplement 2A, 2C-E**). The PCT also exhibited successful proximal migration towards the glomerulus and exhibit the correct convoluted morphology by 3 dpf (**Supplement 2G)**. The observed composition changes were notable as early as the 20 ss and were not a result of changes in cell proliferation, cell death, or total cell number (**Figure 1F, 1G, Supplement 2H-O**). This supports the notion that *esrrγa* operates early specifically throughout nephron formation, as distinct segments are altered.

To further explore the mechanics of these segment changes, we studied another cell type present within the nephron, MCCs. Preceding work from our lab has found that increased monociliated transporter cell identity can be associated with a coordinated decrease in MCC identity (Chambers et al., 2020, Marra et al., 2019a, Marra et al., 2019b). Indeed, we found that *esrrγa* deficiency resulted in a decrease in MCC cell number (marked by *odf3b*), and co-injection of *esrrγa* RNA was sufficient to rescue the splice blocking morpholino (**Figure 1H-I**). Fluorescent in situ hybridization (FISH) analysis of the PST domain (marked by the boundaries of *trpm7*) also revealed a shift towards a transporter cell identity (**Figure 1J**). While the overall average cell number (calculated by number of DAPI) in this domain did not change, there was an increase in the number of *trpm7* positive cells accompanied by a coordinated decrease of *odf3b* positive cells (**Figure 1K**). MCC precursors were also affected in *esrrγa* deficient animals, resulting in a significant decrease in the number of *jag2b* expressing cells (**Supplement 2B, F**). Next, we interrogated *esrrγa* loss of function using a genome editing approach. We designed an independent genetic model of *esrrγa* deficiency using CRISPR-Cas9 mutagenesis. Wild-type zebrafish embryos injected with a cocktail of two guide RNAs that targeted exon 1 of *esrrγa* also exhibited a decrease in MCCs (**Supplement 1G**). As the penetrance of the *esrrγa* crispant phenotypes in F0 mosaic embryos was less consistent than our morpholino models, we continued to use the *esrrγa* morphant models for subsequent analysis. These findings suggest that *esrrγa* contributes to cell fate decisions between the MCC and monociliated transporter cell identity.

### *esrrγa* is required for ciliogenesis in the kidney and other tissues

Ciliogenesis is a complex process, requiring proper basal body production around the centrioles, amplification of the basal bodies (in the case of MCCs), proper basal body docking at the apical surface, and finally cilia outgrowth, mediated by anterograde and retrograde intraflagellar transport (Spassky & Meunier, 2017). Previous studies have found that decreased MCC number can be associated with aberrations in ciliogenesis (e.g. decreased cilia outgrowth) (Chambers et al., 2020).

Considering the observed decrease in MCC number in *esrrγa* deficient animals, we investigated whether cilia formation was likewise affected. The intermediate pronephros are comprised of MCCs interspersed amongst monociliated transporter cells (**Figure 2A**). We first analyzed cryosections of the intermediate nephron of *esrrγa* deficient animals to determine if cell polarity or ciliogenesis was disrupted. We found that morphants successfully established apical basal polarity at 28 hpf, as both apical (aPKC) and basolateral (N+K+ ATPase) proteins were correctly and separately localized (**Figure 2B**). Cryosections of 24 hpf morphants analyzed for cilia structures (cilia (α-tubulin), basal bodies (γ-tubulin)) revealed that while cilia appear to be decreased, basal bodies were successfully docked, as they localized to the putative apical surface (**Figure 2B**). Interestingly, some basal bodies observed in *esrrγa* deficient animals did not appear to be associated with a cilium projection (**Figure 2B**). These data suggest that *esrrγa* could be contributing to proper cilia formation, though this is likely to occur independently of polarity or basal body docking.

**Figure 2.**
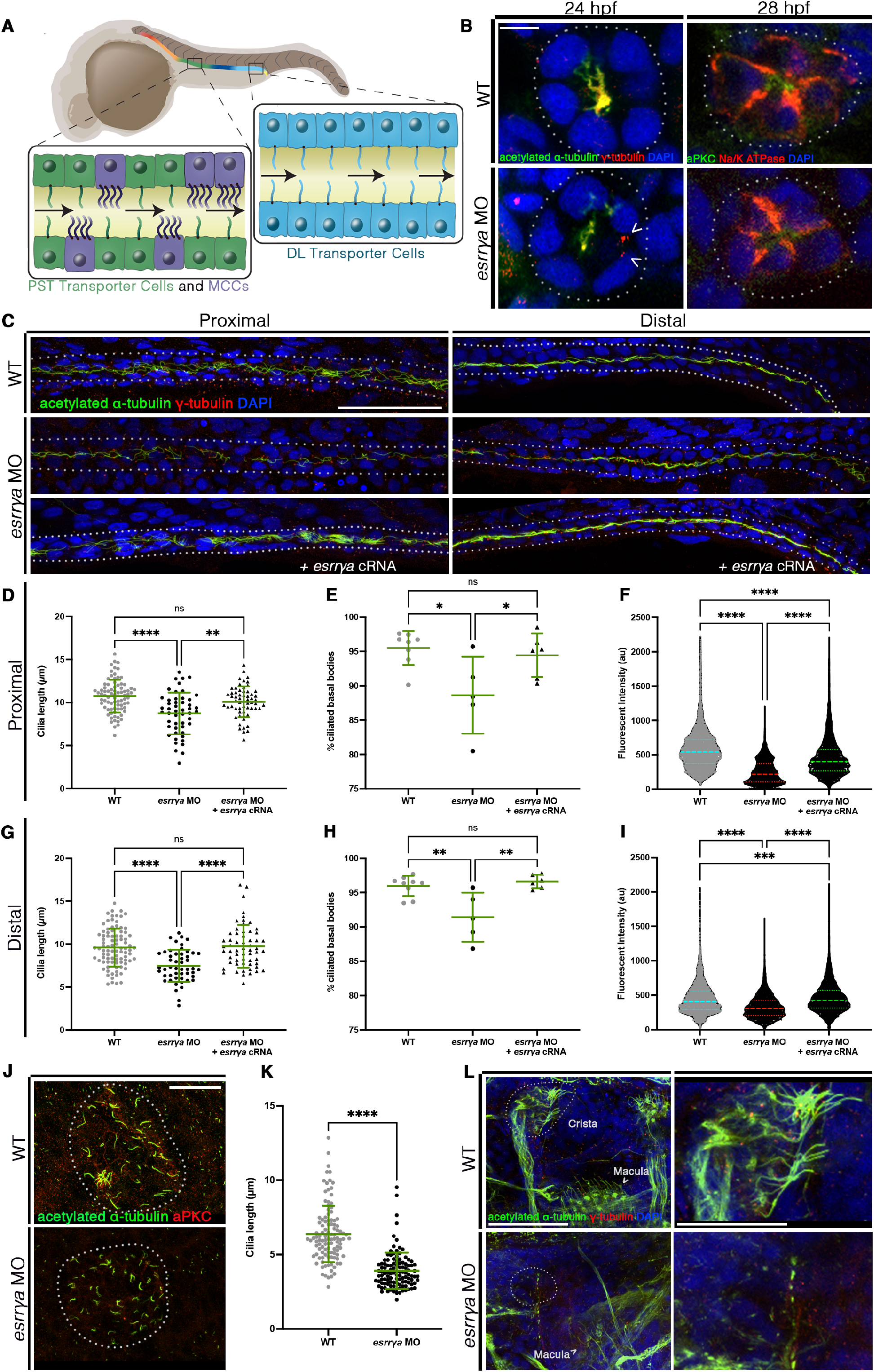
*Esrrγa* contributes to ciliogenesis independent from cell polarity. (A) Schematic illustrating the location of MCCs interspersed amongst monociliated transporter cells in the proximal straight tubule (PST) and monociliated cells along in the distal late (DL) segment. (B) Representative cryosections of the nephron (outlined in dotted line) of WT (top) and *esrrγa* MO injected zebrafish at 24 hpf (left) and 28 hpf (right). Left panels mark cilia (acetylated a-tubulin), and basal bodies (g-tubulin) and. Arrowhead denotes basal body without colocalized cilia. Right panels mark apical surface (aPKC) and basal bodies (Na/K ATPase). Scale bar = 10µm. (C) 28 hpf WT (top), *esrrγa* MO injected (middle), and *esrrγa* MO with *esrrγa* cRNA co-injected (bottom) zebrafish stained via whole mount immunohistochemistry for acetylated a-tubulin (cilia, green), g-tubulin (basal bodies, red), and DAPI in the proximal (left) and distal (right) pronephros. Scale bar = 50µm. (D) Cilia length in micrometers for the proximal pronephros. (E) Percentage of ciliated basal bodies (ciliated basal bodies/total basal bodies per 100µm) in the proximal pronephros. (F) Fluorescent intensity plots (cilia, alpha tubulin) for the same relative distance in the proximal pronephros of WT, *esrrγa* MO, and *esrrγa* MO with *esrrγa* cRNA co-injected animals at 28 hpf. (G) Cilia length in micrometers for the distal pronephros. (H) Percentage of ciliated basal bodies (ciliated basal bodies/total basal bodies per 100µm) in the distal pronephros. (I) Fluorescent intensity plots (cilia, alpha tubulin) for the same relative distance in the distal pronephros of WT, *esrrγa* MO, and *esrrγa* MO with *esrrγa* cRNA co-injected animals at 28 hpf. (J) Representative image of the Kupffer’s vesicle (KV, outlined in dotted line) at the 10 ss stained via immunohistochemistry for acetylated a- tubulin (cilia, green) and aPKC (apical surface, red) for WT (top) and *esrrγa* MO (bottom). Scale bar = 25µm. (K) Cilia length in micrometers for cilia in the KV. (L) Ear structures of WT (top) and *esrrγa* MO (bottom) zebrafish at 4 dpf stained via immunohistochemistry for cilia (green, acetylated alpha tubulin) and basal bodies (red, gamma tubulin). Arrow head denotes macula cilia, dotted line surrounds cristae structures. Dotted area is magnified in right panel. Scale bar left = 50µm, scale bar right = 25µm. Data presented on graphs are represented as mean ± SD; * p<0.05, ** p< 0.01 ***p < 0.001 and ****p < 0.0001 (t test or ANOVA).

To further explore the role of *esrrγa* in cilia outgrowth, we used immunofluorescence to mark cilia (α-tubulin), basal bodies (γ-tubulin), and DAPI in whole mounts of *esrrγa* deficient animals and WT siblings. This was followed by confocal imaging of both the proximal and distal pronephros, to capture cilia protruding from MCCs as well as transporter cells, respectively. Cilia were disrupted in both the proximal and distal pronephros (**Figure 2C, Supplement 3D**), where cilia length was significantly shorter in *esrrγa* deficient animals compared to WT (**Figure 2D, G, Supplement 3E-N**). In addition, morphants did not show any significant changes in the number of basal bodies or cell number (**Supplement 4A-D**). However, *esrrγa* deficient animals had fewer ciliated basal bodies (**Figure 2E, H**), as well as decreased fluorescent intensity of cilia (α-tubulin) (**Figure 2F, I**) which is consistent with the aberrant cilia phenotypes we observed in cryosectioned animals. To test the specificity of our splice interfering morpholino, we co-injected animals with *esrrγa* cRNA. Supplementation of mature *esrrγa* transcript alongside the morpholino was sufficient to rescue cilia length, ciliated basal bodies, and the fluorescent intensity of α-tubulin (**Figure 2C-I**). From these data, we concluded that *esrrγa* deficiency interferes with cilia formation in both MCCs and renal epithelial cells with a single primary cilium.

In addition to the kidney, cilia are critical to several other tissues across vertebrates. In the zebrafish this includes, but is not limited to, the Kupffer’s vesicle (analogous to the mammalian node) and the otic vesicle (ear structure). To determine if *esrrγa* operates solely in the pronephros, we next investigated the effect of *esrrγa* deficiency on these other tissues. We used IF to mark the KV using aPKC (apical surface) and α-tubulin (cilia) in both *esrrγa* morphants and WT siblings at the 10 ss (**Figure 2J**). Like the pronephros, cilia length was significantly reduced in the KV of *esrrγa* deficient animals (**Figure 2K**). We also used IF to identify cilia and basal bodies in the ear at 4 dpf. Similarly, morphant animals exhibited decreased fluorescence of α-tubulin in the region of both macula and cristae structures, the latter of which was nearly absent altogether (**Figure 2L**). These data are consistent with the phenotypes observed in the pronephros, and suggest that *esrrγa* affects multiple tissues throughout development (e.g. 10 ss, 24 hpf, 28 hpf, and 4 dpf).

### *esrrγa* promote ciliogenesis and MCC cell fate by regulating prostanoid biosynthesis

Previous studies from our lab and others have shown that prostaglandin signaling is required for ciliogenesis and MCC cell fate choice (Chambers et al., 2020, Jin et al., 2014, Marra et al., 2019a, Spassky & Meunier, 2017). Considering *esrrγa* deficient animals exhibit similar phenotypes as those with defective prostaglandin synthesis (e.g. decreased MCCs and aberrant cilia), we hypothesized that perhaps *esrrγa* operated in a similar manner. Prostaglandin E_2_ (PGE_2_) was of interest, considering an analog with an improved half-life, 16,16-dimethyl-PGE_2_ (dmPGE_2_), has been able to rescue other animals with aberrant cilia and decreased MCCs (Chambers et al., 2020, Jin et al., 2014, Marra et al., 2019a, Poureetezadi et al., 2016). Using a commercially available ELISA assay, we measured endogenous PGE_2_ levels in WT and *esrrγa* deficient embryos at the 28 hpf stage. Compared to WT, *esrrγa* knockdown resulted in a significant decrease of PGE_2_ (**Figure 3A**). This led us to hypothesize that the diminished PGE_2_ level was the basis for the ciliary and cell fate alterations in *esrrγa* deficient embryos.

**Figure 3.**
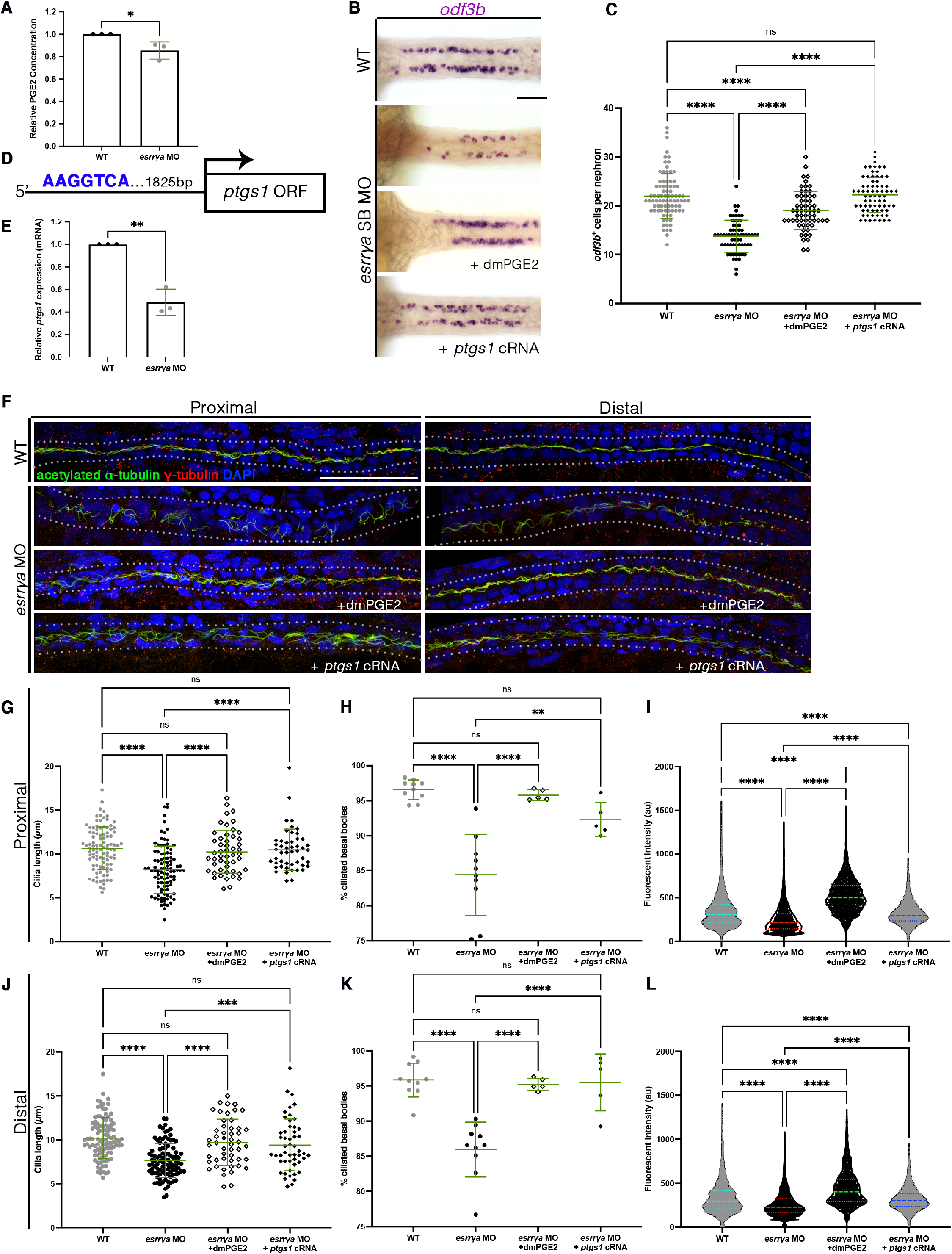
*Esrrγa* controls ciliogenisis through regulation of prostaglandin signaling. (A) Relative PGE2 concentration of 28 hpf WT and *esrrγa* MO injected animals determined via ELISA assay. (B) Representative MCCs stained via WISH (*odf3b*) of 28 ss WT, *esrrγa* MO, *esrrγa* MO treated with dmPGE2, and *esrrγa* MO co-injected with *ptgs1* cRNA (top to bottom, respectively). Scale bar = 50µm. (C) Number of MCCs per nephron of *esrrγa* MO, *esrrγa* MO treated with dmPGE2, and *esrrγa* MO co-injected with *ptgs1* cRNA animals. (D) Schematic illustrating the location of a putative Esrrγa binding site with DNA sequence AAGGTCA upstream of the *ptgs1* open reading frame. (E) Relative amount of *ptgs1* mRNA in WT and *esrrγa* MO animals quantified via qRT-PCR. (F) 28 hpf WT, *esrrγa* MO, *esrrγa* MO treated with dmPGE2, and *esrrγa* MO co-injected with *ptgs1* cRNA (top to bottom, respectively) zebrafish stained via whole mount immunohistochemistry for acetylated a-tubulin (cilia, green), g-tubulin (basal bodies, red), and DAPI in the proximal (left) and distal (right) pronephros. Scale bar = 50µm. (G) Cilia length in micrometers for the proximal pronephros. (H) Percentage of ciliated basal bodies (ciliated basal bodies/total basal bodies per 100µm) in the proximal pronephros. (I) Fluorescent intensity plots (cilia, alpha tubulin) for the same relative distance in the proximal pronephros at 28 hpf. (J) Cilia length in micrometers for the distal pronephros. (K)Percentage of ciliated basal bodies (ciliated basal bodies/total basal bodies per 100µm) in the distal pronephros. (L) Fluorescent intensity plots (cilia, alpha tubulin) for the same relative distance in the distal pronephros at 28 hpf. Data presented on graphs are represented as mean ± SD; * p<0.05, ** p< 0.01 ***p < 0.001 and ****p < 0.0001 (t test or ANOVA).

To test this hypothesis, we examined the functional consequence of restoring prostanoid levels in in *esrrγa* deficient embryos. We treated WT and *esrrγa* deficient embryos with dmPGE_2_ and used WISH of *odf3b* to assess MCC cell fate (**Figure 3B**). *esrrγa* deficient embryos treated with 100 µM dmPGE_2_ from shield stage until fixation at 24 hpf had an increase in MCC number compared to their DMSO treated siblings, restoring the number similar to that of WT animals (**Figure 3C**). As observed by previous studies, we found that dmPGE_2_ was not sufficient to increase MCC number in WT animals (data not shown) (Chambers et al., 2020, Marra et al., 2019a).

Next, we investigated if dmPGE2 was able to restore proper cilia formation. We treated both WT and *esrrγa* morphant animals with 100µM dmPGE2 or vehicle control from shield stage until fixation at 28 hpf and assessed cilia structures using whole mount IF (**Figure 3F**). In both the proximal and distal tubule, dmPGE_2_ rescued cilia length (**Figure 3G, J**), ciliated basal bodies (**Figure 3H, K**), and corresponding cilia fluorescent intensity (**Figure 3I, L**) to WT levels. Together, these data suggest that *esrrγa* interacts with the PGE2 pathway to facilitate ciliogenesis and MCC cell fate.

Prostaglandins are formed by the metabolism of arachidonic acid by cyclooxygenase enzymes to form PGH_2_, which can then be further metabolized by prostaglandin synthase enzymes to form prostanoids. Zebrafish embryos contain four prostaglandin signaling molecules (PGE_2_, PGF2α, PGI_2_, and TXA2) (Cha et al., 2005). Previous research has shown that cyclooxygenase enzymes (Cox1 or Cox2, encoded by *ptgs1, ptgs2a/b* in zebrafish, respectively) are critical for proper ciliogenesis and MCC cell fate (Chambers et al., 2020, Marra et al., 2019a). With this in mind, we hypothesized that *esrrγa* may be contributing to cilia formation through *ptgs1*. We first examined the 2kb promoter region of *ptgs1* for potential binding sites for *esrrγa.* We found one ERR consensus binding motif (AAGGTCA) approximately 1.8kb upstream of the *ptgs1* open reading frame (**Figure 3D**). It is also worth noting that unlike estrogen receptors, ERRs can bind DNA and affect transcription as monomers; thus, one consensus sequence can be sufficient to affect expression (Huppunen et al., 2004). To confirm that *esrrγa* deficiency does affect *ptgs1* transcription, we conducted WISH analysis and qRT-PCR of *ptgs1* in *esrrγa* deficient animals at 24hpf. The length of the *ptgs1* domain in the pronephros was significantly decreased (**Supplement 5A-B**), as well as expression in whole animal lysates, as determined by qRT-PCR (**Figure 3E**). The observed decreased expression of *ptgs1* further supports our hypothesis *esrrγa* may be contributing to cilia formation through regulation of cyclooxygenase enzyme 1. Next, we sought to determine if *ptgs1* overexpression alone was sufficient to rescue *esrrγa* deficiency. We first examined the effect of co-injection of *ptgs1* RNA with *esrrγa* MO on MCC number (**Figure 3B**). Like dmPGE_2_ treatment, *ptgs1* RNA restored MCC number to WT levels (**Figure 3C**). We then observed the effect of *ptgs1* overexpression on cilia in both MCCs and mono-ciliated cells using IF (**Figure 3F**). In both proximal and distal segments, we found that *ptgs1* cRNA was sufficient to rescue cilia length (**Figure 3G, J**), ciliated basal bodies (**Figure 3H, K**), and the corresponding fluorescent intensity of α-tubulin (**Figure 3I, L**). Again, these phenotypes were not due to changes in basal body or cell number (**Supplement 4A-D**). From these data, we concluded that *esrrγa* promotes PGE_2_ synthesis via *ptgs1* to promote MCC specification and cilia outgrowth in both MCC and transporter cell populations.

### *esrrγa* cooperates with *ppargc1a* to control MCC specification and cilia formation

Recent studies have found that *ppargc1a* is essential for prostaglandin signaling, nephron formation, and ciliogenesis (Chambers et al., 2018, Chambers et al., 2020). Interestingly, deficiency of this factor results in similar phenotypes that we observed in the case of *esrrγa* deficiency, including decreased DL and skewed MCC cell fate choice (Chambers et al., 2018, Chambers et al., 2020). Considering these similarities, we sought to determine if *esrrγa* and *ppargc1a* are expressed in the same cell population. FISH analysis revealed that *esrrγa* and *ppargc1a* colocalize in the same pan-distal region of the nephron (**Figure 4A**).

**Figure 4.**
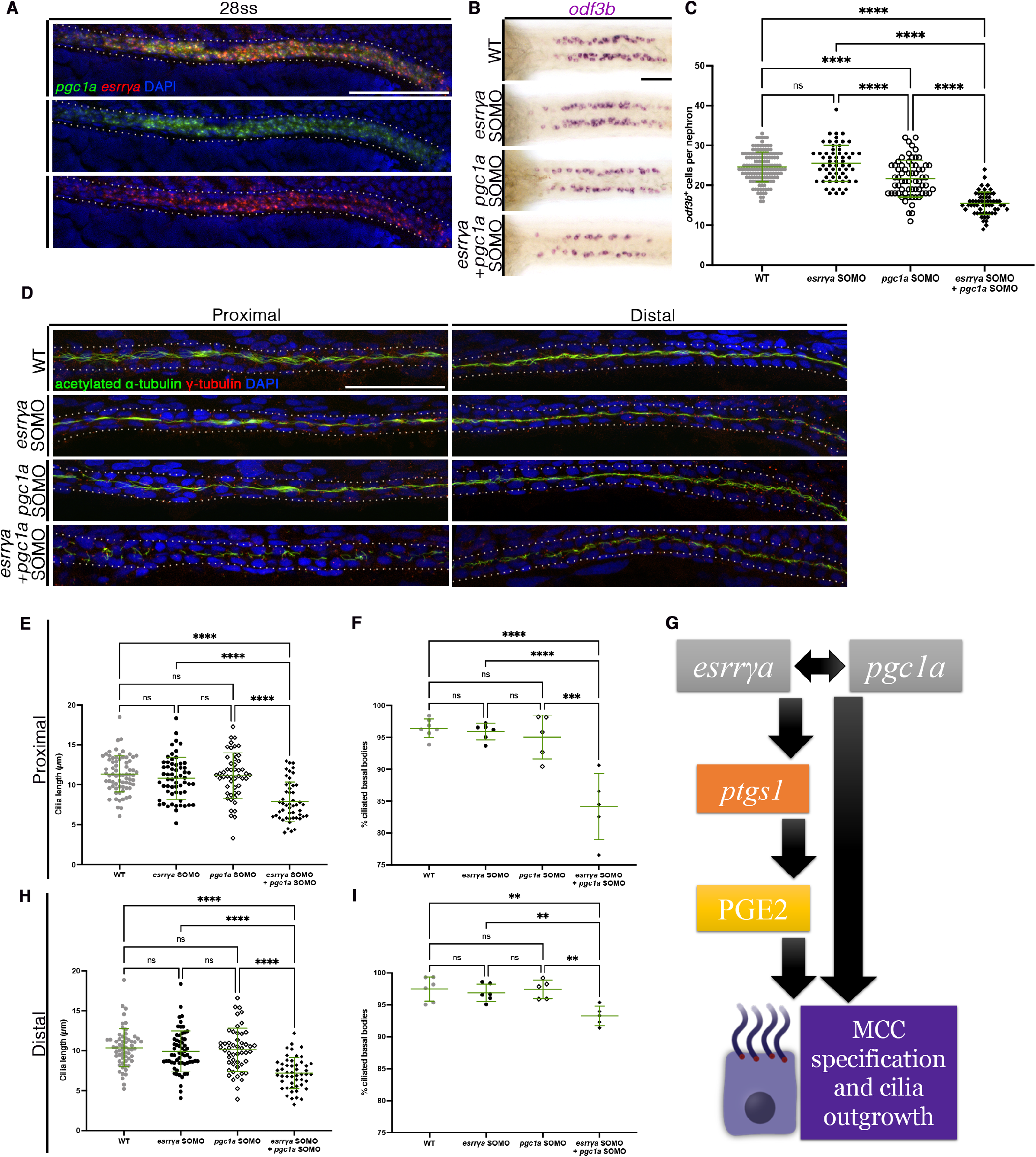
*Esrrγa* cooperates with *pgc1a* to facilitate MCC specification and cilia outgrowth. (A) Expression of *pgc1a* (green) and *esrrγa* (red) marked in fluorescent *in situ* hybridization in a 24 hpf WT zebrafish Scale bar = 50µm. (B) Representative MCCs stained via WISH (*odf3b*) of 24 ss WT, *esrrγa* sub-optimal MO (SOMO), *pgc1a* SOMO, and *esrrγa* SOMO and *pgc1a* SOMO co- injected (top to bottom, respectively). Scale bar = 50µm. (C) Number of MCCs per nephron. (D) 28 hpf WT, *esrrγa* sub-optimal MO (SOMO), *pgc1a* SOMO, and *esrrγa* SOMO and *pgc1a* SOMO co- injected (top to bottom, respectively) zebrafish stained via whole mount immunohistochemistry for acetylated a-tubulin (cilia, green), g-tubulin (basal bodies, red), and DAPI in the proximal (left) and distal (right) pronephros. Scale bar = 50µm. (E) Cilia length in micrometers for the proximal pronephros. (F) Percentage of ciliated basal bodies (ciliated basal bodies/total basal bodies per 100µm) in the proximal pronephros. (G) Model illustrating MCC cell fate and cilia outgrowth by which Esrrγa and Pgc1a cooperate upstream of *ptgs1* and PGE2 production. (H) Cilia length in micrometers for the distal pronephros. (I) Percentage of ciliated basal bodies (ciliated basal bodies/total basal bodies per 100µm) in the distal pronephros. Data presented on graphs are represented as mean ± SD; * p<0.05, ** p< 0.01 ***p < 0.001 and ****p < 0.0001 (t test or ANOVA).

Considering these factors are co-expressed, we designed genetic interaction studies to explore the relationship between *esrrγa* and *ppargc1a*. One strategy to test if multiple genes operate synergistically in a pathway is the use of suboptimal morpholinos (Chambers et al., 2020, Choi et al., 2015, DiBella et al., 2009, Kallakuri et al., 2015, Wagle et al., 2011). Therefore, we injected suboptimal morpholino (SOMO) doses of both *esrrγa* and *ppargc1a*, based on previously published doses (Chambers et al., 2018, Chambers et al., 2020). We then conducted WISH to determine changes in MCC number (**Figure 4B**). We found that *esrrγa* and *ppargc1a* SOMO alone resulted in no change or a slight yet significant decrease in MCC number when compared to WT, respectively (**Figure 4C)**. However, the combination of both *esrrγa* and *ppargc1a* SOMO together resulted in a significant decrease in MCC number (**Figure 4C**).

We then interrogated the synergistic effect of *esrrγa* and *ppargc1a* on cilia formation. Similar to MCC number, *esrrγa* and *ppargc1a* SOMO did not appear to have a significant effect on the appearance of cilia, while the combination injection showed aberrant cilia structures (**Figure 4D**). Further, *esrrγa* and *ppargc1a* SOMO injections independently did not significantly change ciliated basal bodies nor cilia length in MCC and transporter cell populations. However, combination of the *esrrγa* and *ppargc1a* SOMO significantly decreased the percentage of ciliated basal bodies as well as cilia length in both pronephric regions of interest (**Figure 4E-F, H-I**). The corresponding fluorescent intensity of the combination SOMO was also significantly decreased when compared to all other treatment groups (**Figure 4G, 4J**). These changes were not due to alterations in basal body or cell number (**Supplement 4I-L**). Overall, this evidence is indicative of a cooperative effect between *esrrγa* and *ppargc1a* in the context of ciliogenesis and MCC specification.

## Discussion

Estrogen related receptors, and specifically Esrrγ, have been previously linked to disease states of tissues with high energy demand. Global Esrrγ knockout mice and cardiac specific overexpression mice exhibit early lethality due to heart failure (Alaynick et al. 2007, Alaynick et al. 2010, Lasheras et al., 2021). Similar dysfunction is seen in the kidney, as Esrrγ knockout results in deficient ureteric branching, kidney cysts, and decreased mitochondrial function and solute transportation (Berry et al., 2011, Zhao et al., 2018). In humans, mutations in or decreased expression of ESRRγ has been linked to incidence of congenital anomalies of the kidney and urinary tract and chronic kidney disease (Berry et al., 2011, Eichner et al., 2011, Hock et al., 2009, Misra et al., 2017, Zhao et al., 2018). Yet, until our current work, the mechanism by which Esrrγ contributes to the development of the high energy and ciliated tissues remains poorly understood.

Our work suggests that *esrrγa* works with *ppargc1a* upstream of prostaglandin signaling to facilitate nephron cell development and ciliogenesis. Specifically, *esrrγa* acts as a “switch” to favor MCC fate, as we see decreased MCC number with a coordinated increase of PST transporter cells and decrease of the DL segment in *esrrγa* deficiency. Additionally, *esrrγa* knockdown resulted in decreased ciliated basal bodies and decreased cilia length in both mono and multiciliated cell populations in the nephron and other tissues. The observed decreased MCC number and aberrant cilia could be rescued by co-injection of *ptgs1* or treatment with dmPGE_2_, showing for the first time that *esrrγa* works upstream of prostanoid production. Furthermore, suboptimal morpholino injection of *esrrγa* and *ppargc1a*, genetically mimicking compound heterozygous animals, resulted in phenotypes reminiscent of full dosage animals with decreased MCC number and atypical cilia. Together, these data deepen our understanding of the possible mechanisms contributing to ciliopathic and kidney disease conditions.

Prior to this work, prostaglandins have been established as key bioactive molecule in various tissues, and implicated in disease states relating to inflammation, vascular development, cardiac injury, and kidney disease (Lannoy et al., 2020, FitzSimons et al., 2020, Marra et al., 2019a, Sparks and Coffman, 2010, Ugwuagbo et al., 2019,). In some models, blockade of a prostaglandin receptor (EP4) can improve cystic disease states, yet PGE_2_ has also been implicated as an essential factor of cilia outgrowth and proper nephron patterning, which together point to the importance of proper spatiotemporal control of prostaglandin dosage. (Chambers et al., 2020, Jin et al., 2017, Lannoy et al., 2020, Marra et al., 2019a, Poureetezadi et al., 2016). Here, we have added to that growing body of knowledge, as dmPGE2 and *ptgs1* were able to rescue both cell type (MCC deficiency) and ciliopathic (ciliated basal bodies, and cilia length) phenotypes. The dmPGE2 treatment was unable to rescue the distal late segment length (data not shown). However, a restoration of the DL was not anticipated as exogenous dmPGE_2_ and prostaglandin inhibition have been shown to result in the same DL domain decrease (Poureetezadi et al., 2016). These somewhat contradictory findings speak to the importance of precise PGE_2_ dosage, and also suggests that *esrrγa* may control segmentation of the distal segment through a mechanism independent of prostaglandin signaling. Further research is required to interrogate the mechanism by which the distal segment is regulated. Candidate transcription factors like *tbx2b* may be of interest, as it operates downstream of *ppargc1a* in distal cell fate identity (Chambers et al., 2018, Drummond et al., 2017).

*esrrγa* and *ppargc1a* act similarly upstream of nephron and cilia development in zebrafish, and we found a synergistic relationship these factors. Ciliogenesis is not the first context in which these factors have been linked, and some have even suggested that PGC1a acts as a “protein ligand” of ESRRγ (Audet-Walsh and Giguere, 2014). Both *esrrγa* and *ppargc1a* have been independently implicated in mitochondrial function and various disorders, including diabetes and kidney disease (Audet-Walsh and Giguere, 2014, Guo et al., 2015, Ishimoto et al., 2017, Knutti and Kralli, 2001, Long et al., 2016, Misra et al., 2017, Poidatz et al., 2012, Sharma et al., 2013, Zhao et al., 2018). Further, Esrrγ and Ppargc1a have been shown to bind the same multiple hormone response element in the context of kidney specific and other cell lines (Liu et al., 2005, Wang et al., 2008). While the present studies have shown a strong and consistent interaction between *esrrγa* and *ppargc1a*, cilia structure and specific nephron cell types were not evaluated. While we have begun filling this gap in knowledge through our suboptimal morpholino combination experiments, future studies are needed to elucidate the molecular nature of this relationship. In particular, it is not yet known if Esrrγa and Ppargc1a directly bind to one another in the promoter or enhancer region of *ptgs1* or regulate ciliogenesis through some other mechanism. Since both Esrrγ and Ppargc1a have been shown to interact with Hnf1b in the context of kidney tissue, especially in ciliopathic conditions, it is possible that this factor may act as the link in this synergistic relationship (Casemayou et al., 2017, Verhave et al., 2016, Zhao et al., 2018). Furthermore, neither *esrrγa* or *ppargc1a* deficiency alone is sufficient to eradicate all pronephric MCCs and cilia. This may be due to redundant function of *esrrγb*, as *esrrγa/b* function redundantly in the development of the otic vesicle (Tohme et al., 2014). Alternatively, maternal deposition of either of these factors could explain the basal level of cilia production or perhaps the presence of other ciliogenic factors is sufficient to compensate for *esrrγa* and *ppargc1a* loss to drive low levels of ciliogenesis. Future studies may be interested in the interaction with *esrrγa/ppargc1a* and other known components of the ciliogenesis network (e.g. *foxj1*, *gmnc,* and *mulitcillin*) (Choksi et al., 2014, Spassky and Meunier, 2017). It is possible that complete absence of MCCs and cilia is only possible when animals are deficient in multiple factors within the cilia regulatory program.

The link between ciliopathies and aberrant kidney structure and function has long been established (Wang et al., 2013, Winyard et al., 2011), yet the relationship between nephrogenesis and ciliogenesis remains poorly understood. Our work has identified a novel role for *esrrγa* at the nexus of kidney and cilia formation through prostaglandin signaling and cooperation with *ppargc1a*. This discovery suggests that Esrrγ is an important component in maintaining kidney health and implicates Esrrγ as a key regulator of ciliogenesis in other tissues. Other studies have already recognized the potential of ERRs as targets for aging kidney treatment. Specifically, treating with pan-ERR agonists results in remarkable improvements in mitochondrial function and albuminaria (Wang et al., 2020). Our findings support this trend, as Esrrγ may serve as a future therapeutic target, and considering the worldwide prevalence of kidney failure and ciliopathies, novel targets are of the upmost importance.

## ACKNOWLEDGEMENTS

We would like to thank the generous funders of this work: startup funds from the University of Notre Dame (to R.A.W), Graduate Women in Science National Fellowship (to H.M.W), Warren Center Drug Development Welter Family Fellowship (to H.M.W), and the Notre Dame Center for Stem Cells and Regenerative Medicine Fellowship (to H.M.W). This work would not have been possible without the staffs of the Department of Biological Sciences and the Center for Zebrafish Research at the University of Notre Dame. Imaging seen in this manuscript was carried out in part in the Notre Dame Integrated Imaging Facility (AR1, C2 confocal microscopes), and we especially thank S.C. for her knowledge and expertise. Finally, we express our deep gratitude to the Wingert lab and Wingert lab alumni, J.C and B.C., for their guidance and insight on this project.

## Methods

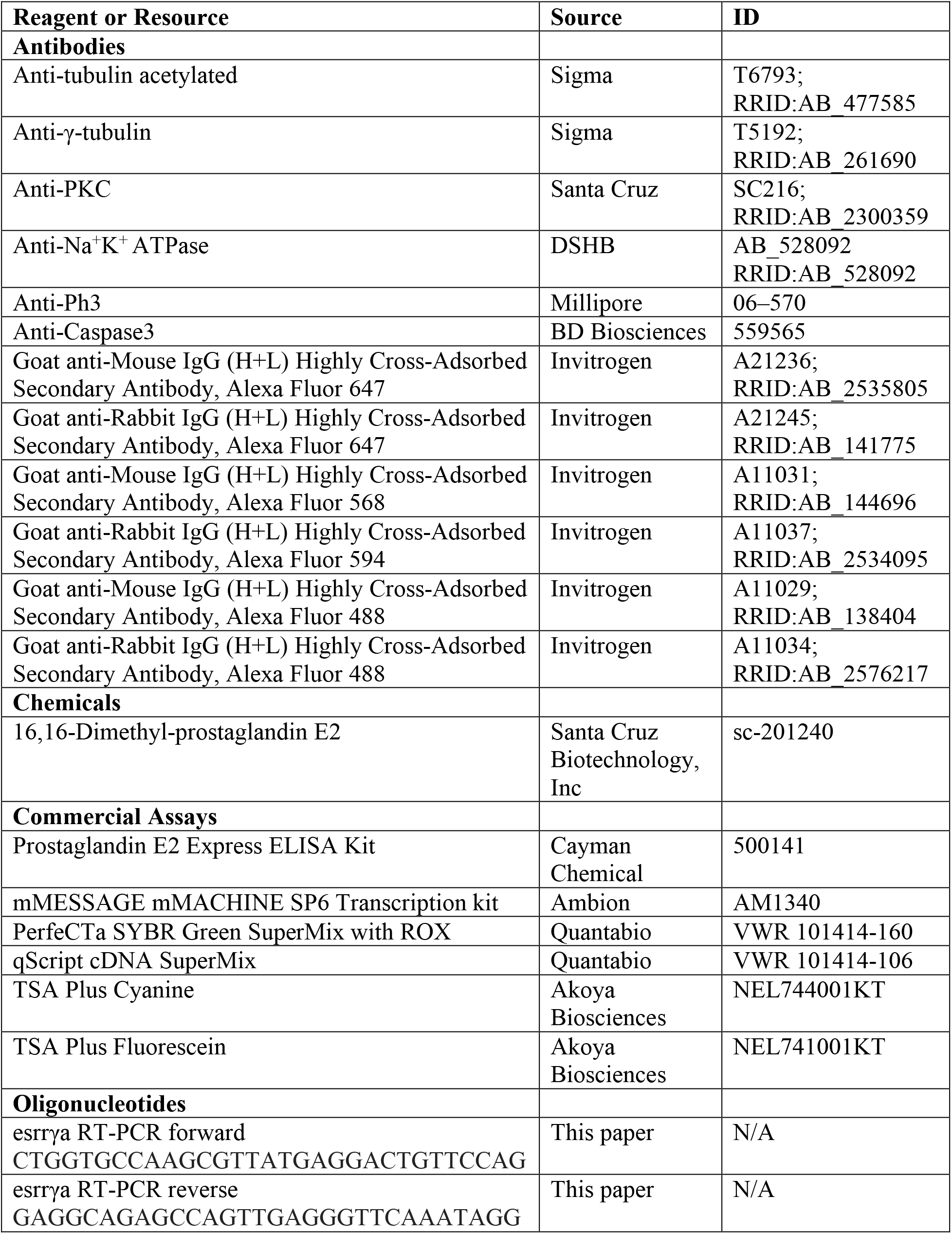
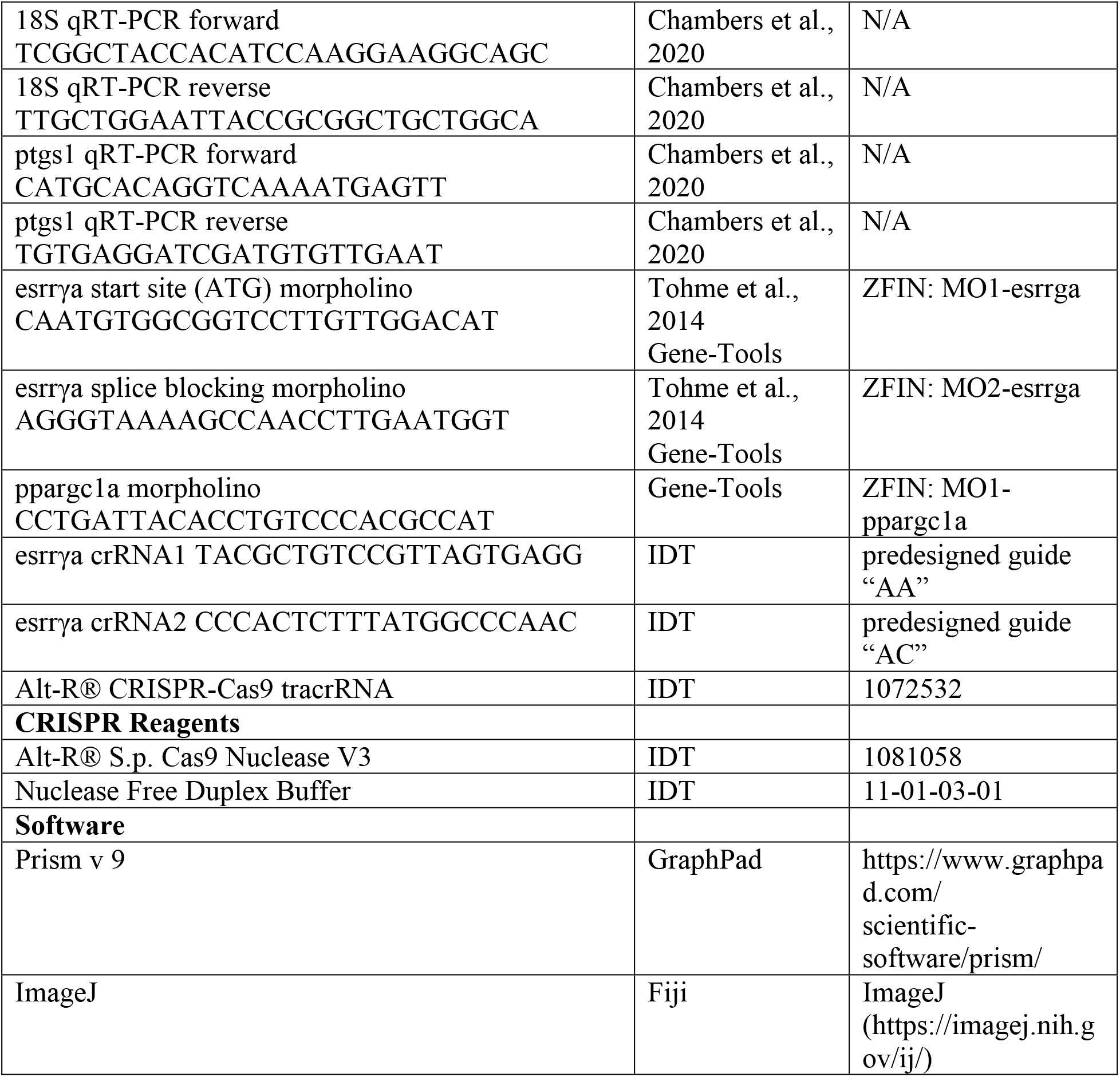

### EXPERIMENTAL MODEL AND SUBJECT DETAILS

The Center for Zebrafish Research at the University of Notre Dame maintained the zebrafish used in these studies and experiments were performed with approval of the University of Notre Dame Institutional Animal Care and Use Committee (IACUC), under protocol number 19-06-5412.

#### Animal models

Tübingen strain WT zebrafish were used for all studies. Zebrafish were raised and staged as described (Kimmel et al., 1995). For all studies, embryos were incubated in E3 medium at 28°C until the desired developmental stage, anesthetized with 0.02% tricaine, and then fixed using 4% paraformaldehyde/1x PBS (PFA), or Dent’s solution (80% methanol, 20% DMSO) (Westerfield, 1993, Gerlach et al., 2014). Embryos were analyzed before sex determination, so we cannot report the effect of sex and gender in the context of this study.

### METHOD DETAILS

#### Whole mount and fluorescent whole mount in situ hybridization (WISH, FISH)

WISH was performed as previously described (Cheng et al., 2014; Galloway et al., 2008; Lengerke et al., 2011; Marra et al., 2019a, Chambers et al 2020) with antisense RNA probes either digoxigenin-labeled (*esrrγa, cdh17, odf3b, slc20a1a, trpm7, slc12a1, slc12a3, jag2b, ptgs1*) or fluorescein-labeled (*deltaC, smyhc, pax2a, odf3b, esrrγa, cdh17, pgc1a, slc12a3*) using *in vitro* transcription using IMAGE clone templates as previously described (Wingert et al., 2007; O’Brien et al., 2011; Gerlach and Wingert, 2014). FISH was performed as described (Brend and Holley 2009; Marra et al., 2017) using TSA Plus Fluorescein or Cyanine Kits (Akoya Biosciences). For all gene expression studies, every analysis was done in triplicate for each genetic model with sample sizes of n > 20 per replicate.

#### Sectioning

Fixed zebrafish samples exposed to 5% and 30% sucrose solution and then subjected to a 1:1 solution of 30% sucrose and tissue freezing medium (TFM). Infiltrated samples were embedded in 100% TFM and oriented in Tissue-Tekcryo-molds and frozen at -80°C. Sections (14 µm) were taken on a Microm HM 550 Cryostat (Thermo).

#### Immunofluorescence (IF)

Whole mount IF experiments were completed as previously described (Gerlach and Wingert, 2014; Kroeger et al., 2017; Marra et al.,2017, 2019c, Chambers et al., 2020). For cilia and basal bodies, anti-tubulin acetylated diluted 1:400 (Sigma T6793) and anti γ-tubulin diluted 1:400 (Sigma T5192) were used, respectively. Cryosectioned samples were completed as previously described (Gerlach and Wingert, 2014). For cilia and basal bodies, anti-tubulin acetylated diluted 1:1000 (Sigma T6793) and anti γ-tubulin diluted 1:400 (Sigma T5192). For cell polarity, animals were fixed in Dent’s solution, and used anti-aPKC diluted 1:500 (Santa Cruz 2300359) to mark apical surface and anti-Na^+^K^+^ ATPase diluted 1:35 (DSHB 528092) for a basolateral marker.

#### Rescue Experiments with dmPGE2

Chemical treatments were completed as previously described (Marra et al., 2019a; Poureetezadi et al., 2014; Poureetezadi et al., 2016, Chambers et al., 2020). 16,16-dmPGE_2_ (Santa Cruz Biotechnology, Inc, SC-201240) was dissolved in 100% dimethyl sulfoxide (DMSO) to make 1M stocks then diluted to the 100 µM treatment dose. Treatments were completed in triplicate with n > 20 embryos per replicate. PGE_2_ metabolite quantification PGE_2_ metabolite quantifications were completed according to the manufacturer’s protocol (Cayman Chemical #500141). In brief, groups of 25 WT or *esrrγa* MO injected zebrafish were pooled, anesthetized, and flash frozen in 100% ethanol. Lysates were homogenized and supernatant was isolated after centrifugation at 4°C (12,000 RPM for 20 minutes). The kit reagents and manufacturer supplied protocol was followed for assay completion using a plate reader (SpectraMax ABSPlus) at 420 nm.

#### Quantitative real-time PCR

Groups of 30 zebrafish (WT, *esrrγa* morphants) were pooled at 24 hpf. Trizol (Ambion) was used to extract RNA, qScript cDNA SuperMix (QuantaBio) was used to make cDNA. PerfeCTa SYBR Green SuperMix with ROX (QuantaBio) was used to complete qRT-PCR with 100 ng for ptgs1 and 1 ng for 18S controls being optimal cDNA concentrations. The AB StepOnePlus qRT-PCR machine was used with the following program: 2 minute 50°C hold, 10 minute 95°C hold, then 35 cycles of 15 s at 95°C and 1 minute at 60°C for denaturing and primer annealing and product extension steps respectively. Each target and source were completed in biological replicates and technical replicates each with the median Ct value normalized to the control. Data analysis was completed by using delta delta Ct values comparing WT uninjected siblings to the respective morphant groups with 18S as a reference. Primers used include: To target 18S: forward 5’–TCGGCTACCACATCCAAGGAAGGCAGC–3’ reverse 5’–TTGCTGGAATTACCGCGGCTGCTGGCA–3’.

To target *ptgs1*: forward 5’- CATGCACAGGTCAAAATGAGTT- 3’ reverse 5’-TGTGAGGATCGATGTGTTGAAT-3’ cRNA synthesis, and microinjections, rescue studies The zebrafish *esrrγa* ORF was cloned in to a pUC57 vector flanked by a 5’ KOZAK sequence, single BamH1, SalI and EcoRV restriction sites, and a SP6 promoter region. On the 3’ side, the ORF was followed by a series of STOP codons, a SV40 poly A tail, single NDeI, EcoRI, and NotI restriction sites, and a T7 promoter region. *esrrγa* RNA was generated by linearizing with Not1 and SP6 run off with the mMESSAGE mMACHINE SP6 Transcription kit (Ambion). *esrrγa* RNA was injected into WT with or without a co-injection of *esrrγa* splice blocking morpholino at the 1- cell stage at a concentration of 500 pg. The *ptgs1* ORF was cloned in to a pUC57 vector flanked by a 5’ KOZAK sequence, Cla1 restriction site, and a SP6 promoter region. On the 3’ side, the ORF was followed by a series of STOP codons, a SV40 poly A tail, a NotI restriction site, and a T7 promoter region. *ptgs1* RNA was generated as with *esrrγa* and injected at 900 pg.

#### CRISPR-Cas9 mutagenesis

Methods were adapted from Hoshijima et al., (2019). In short, target sequences were selected using the IDT predesign tool, and cross referenced using online program CHOP-CHOP (http://chopchop.cbu.uib.no/index.php). Selected crRNA and tracrRNA tools were obtained (IDT), and dissolved into a 100µM stock with duplex buffer (IDT). To form the crRNA:tracrRNA duplex, equal amounts of crRNA were combined with tracrRNA, and exposed to a rapid heat-slow cool program in a thermocycler. The 50µM duplexed crRNA:tracrRNA was diluted to 25µM with duplex buffer (IDT). Cas9 protein (IDT) was prepared and aliquoted according to the protocol described by Hoshijima et al., (2019). Injection mixes were prepared as follows: 1µl 25µM crRNA:tracrRNA (crRNA 1) + 1µl 25µM crRNA:tracrRNA (crRNA 2) + 1µl 25µM Cas9 protein + 2µl RNase-free water. This mixture was incubated at 37°C for 10 minutes, and then stored at room temperature. Zebrafish embryos were injected at the one cell stage with 2-3nl of the 5µM crRNA:tracrRNA:Cas9 mixture.

#### Genetic models

Antisense morpholino oligonucleotides (MOs) were obtained from Gene Tools, LLC (Philomath, OR). MOs were solubilized in DNase/RNase free water to generate 4 mM stock solutions which were stored at 20C. Zebrafish embryos were injected at the 1-cell stage with 1-2 nL of diluted MO. *esrrγa* was targeted with two morpholinos. A start site (ATG) morpholino: 5’– CAATGTGGCGGTCCTTGTTGGACAT –3’ (667 µM optimal), and a splice blocking morpholino 5’– AGGGTAAAAGCCAACCTTGAATGGT –3’(400 µM optimal, 200 µM suboptimal). The latter of which was validated using RT-PCR using the following primers: *esrrγa* RT-PCR forward 5’– CTGGTGCCAAGCGTTATGAGGACTGTTCCAG –3’ and *esrrγa* RT-PCR reverse 5’– GAGGCAGAGCCAGTTGAGGGTTCAAATAGG–3’. *ppargc1a* was targeted with the following validated MO: 5’–CCTGATTACACCTGTCCCACGCCAT–3’ (400 µM optimal, 200 µM suboptimal) (Hanai et al., 2007; Bertrand et al., 2007, Chambers et al., 2018, Chambers et al., 2020).

#### Image acquisition and phenotype quantification

A Nikon Eclipse Ni with a DS-Fi2 camera was used to image WISH samples and live zebrafish. Live zebrafish were mounted in methylcellulose with trace amounts of tricaine present. IF and FISH images were acquired using a Nikon C2 confocal microscope

### QUANTIFICATION AND STATISTICAL ANALYSIS

Cilia phenotypes were quantified using ImageJ/Fiji (https://imagej.nih.gov) software tools. All measurements were completed on representative samples imaged at 60X magnification. The multi-point tool was used for counting. The segmented line tool was used for length measurements. Fluorescent intensity plots were generated with the plot profile function. Each experiment was completed in a minimum of triplicate. From these measurements an average and standard deviation (SD) were calculated, and unpaired t tests or one-way ANOVA tests were completed to compare control and experimental measurements using GraphPad Prism 9 software. Statistical details for each experiment are located in the corresponding figure legend.

## Supplemental Figures and Figure Legends

**Supplement 1.**
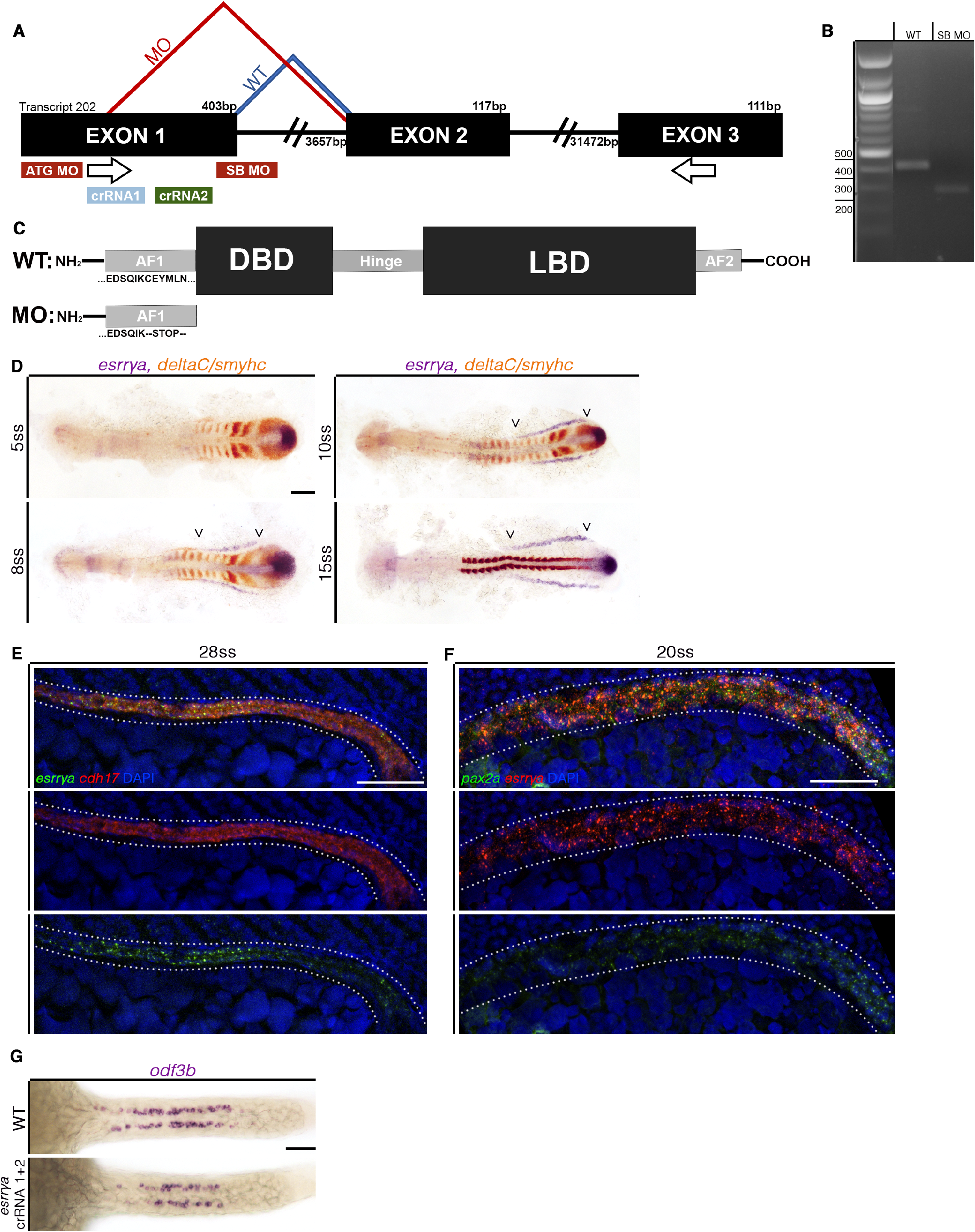
(A) Schematic illustrating morpholino knockdown tools. Start site (ATG) MO binds to the ATG start site in exon 1. Splice blocking (SB) MO binds to the first exon-intron junction. Dark blue lines indicate WT splicing, while red lines indicate altered splicing resulting from splice blocking MO injection. Arrows show primer locations used for RT-PCR. crRNA1 (light blue) and crRNA2 (green) indicate the relative locations of the guide RNAs used for CRISPR-Cas9 mutagenesis. (B) DNA agarose gel of RT-PCR products of WT (Lane 2) and splice blocking MO (Lane 3) animals. WT band is present at the anticipated 450 bp length while the MO band is smaller (approximately 300 bp), indicating that part of exon 1 was spliced out. Bands were gel purified and confirmed with sequencing analysis. (C) Schematic illustrating WT protein domains (top), and putative protein resulting from splice blocking MO injection (bottom). (D) *esrrγa* expression (purple) and somite location (*deltaC/smyhc*, orange) stained via WISH at various stages. Arrow heads indicate *esrrγa* expression domain. Scale bar = 100µm. (E) FISH expression of *esrrγa* (green) and *cdh17* (red) at the 28 ss. Top is the merged file, middle is *cdh17* alone, and bottom is *esrrγa* alone. Scale bar = 50µm. (F) FISH expression of *esrrγa* (red) and *cdh17* (pax2a) at the 20 ss. Top is the merged file, middle is *esrrγa* alone, and bottom is *pax2a* alone. Scale bar = 50µm. (G) Representative image of MCCs stained with *odf3b* via WISH of 24 hpf WT (top) and a confirmed CRISPR-Cas9 *esrrγa* mutant (bottom). Mutant was confirmed via T7 endonuclease assay (data not shown). Scale bar = 50µm.

**Supplement 2.**
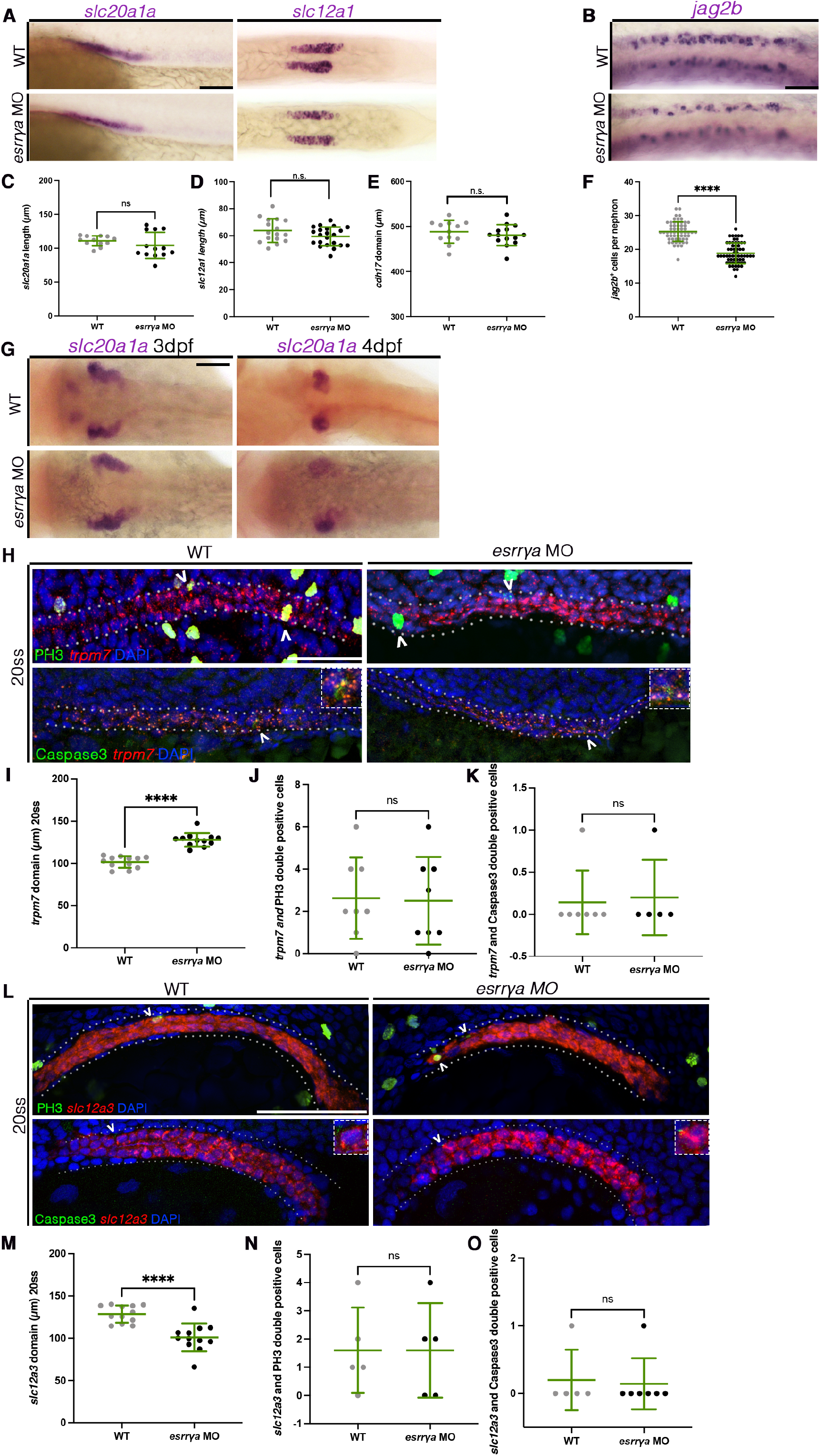
(A) WISH of WT (top) and *esrrγa* MO (bottom) zebrafish at 24 hpf stained for PCT (*slc20a1a*, left), or DE (*slc12a1,* right). Scale bar = 50µm. (B) WISH of WT (top) and *esrrγa* MO (bottom) zebrafish at 24 hpf stained for MCC precursors (*jag2b*). Scale bar = 50µm. (C) PCT domain length in micrometers. (D) DE domain length in micrometers. (E) Entire nephron tubule length (representative image seen in Figure 1D) in micrometers. (F) Number of MCC precursors per nephron. (G) Representative images of the PCT, marked by WISH of *slc20a1a*, during convolution of WT (top) and *esrrγa* MO (bottom) animals and 3dpf (left) and 4 dpf (right). Scale bar = 50µm. (H) 20ss WT (left) and *esrrγa* MO (right) nephrons (outlined with dotted line) stained for the PST (*trpm7*) via fluorescent in situ hybridization and proliferating (PH3, top) or apoptotic (Caspase3, bottom) cells via immunohistochemistry. Arrow heads denote double positive cells. Scale bar = 50µm. (I) PST (*trpm7*) domain length at 20ss in micrometers. (J) Number of PH3 positive cells in the PST at 20 ss (K) Number of Caspase3 positive cells in the PST at 20 ss. (L) 20 ss WT (left) and *esrrγa* MO (right) nephrons (outlined with dotted line) stained for the DL (*slc12a3*) via fluorescent in situ hybridization and proliferating (PH3, top) or apoptotic (Caspase3, bottom) cells via immunohistochemistry. Arrow heads denote double positive cells. Scale bar = 50µm. (M) DL (*slc12a3*) domain length at 20 ss in micrometers. (N) Number of PH3 positive cells in the DL at 20 ss. (O) Number of Caspase3 positive cells in the DL at 20 ss.

**Supplement 3.**
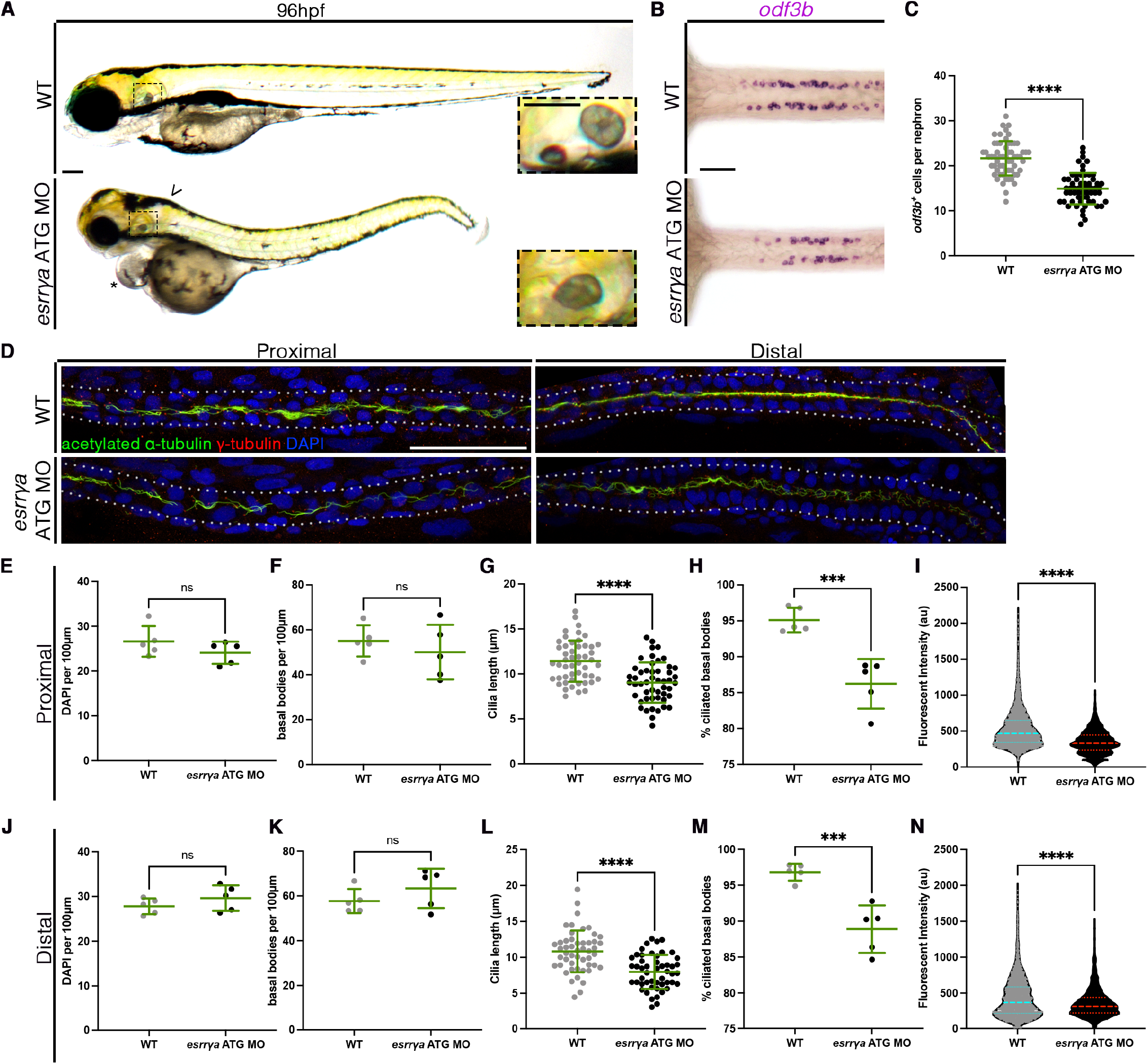
(A) WT sibling (top) and e*srrγa* morphant (bottom) zebrafish with pericardial edema (asterisk), hydrocephaly (arrow head), and fused otoliths (dashed box outline, inset). Scale bar = 100µm, inset = 50µm. (B) WT (top) and *esrrγa* MO injected (bottom) zebrafish at 24 hpf stained with WISH for MCCs (*odf3b*). Scale bar = 50µm. (C) Absolute number of MCCs per nephron of zebrafish at 24 hpf (D) 28 hpf WT (top) and *esrrγa* MO (bottom) zebrafish stained via whole mount immunohistochemistry for acetylated a-tubulin (cilia, green), g-tubulin (basal bodies, red), and DAPI in the proximal (left) and distal (right) pronephros. Scale bar = 50µm. (E) Number of DAPI cells per 100µm in the proximal pronephros. (F) Number of basal bodies per 100µm in the proximal pronephros. (G) Cilia length in micrometers in the proximal pronephros. (H) Percentage of ciliated basal bodies (ciliated basal bodies/total basal bodies per 100µm) in the proximal pronephros. (I) Fluorescent intensity plots (cilia, alpha tubulin) for the same relative distance in the proximal pronephros at 28 hpf. (J) Number of DAPI cells per 100µm in the distal pronephros. (K) Number of basal bodies per 100µm in the distal pronephros. (L) Cilia length in micrometers for the distal pronephros. (M) Percentage of ciliated basal bodies (ciliated basal bodies/total basal bodies per 100µm) in the distal pronephros. (N) Fluorescent intensity plots (cilia, alpha tubulin) for the same relative distance in the distal pronephros at 28 hpf. Data presented on graphs are represented as mean ± SD; * p<0.05, ** p< 0.01 ***p < 0.001 and ****p < 0.0001 (t test or ANOVA).

**Supplement 4.**
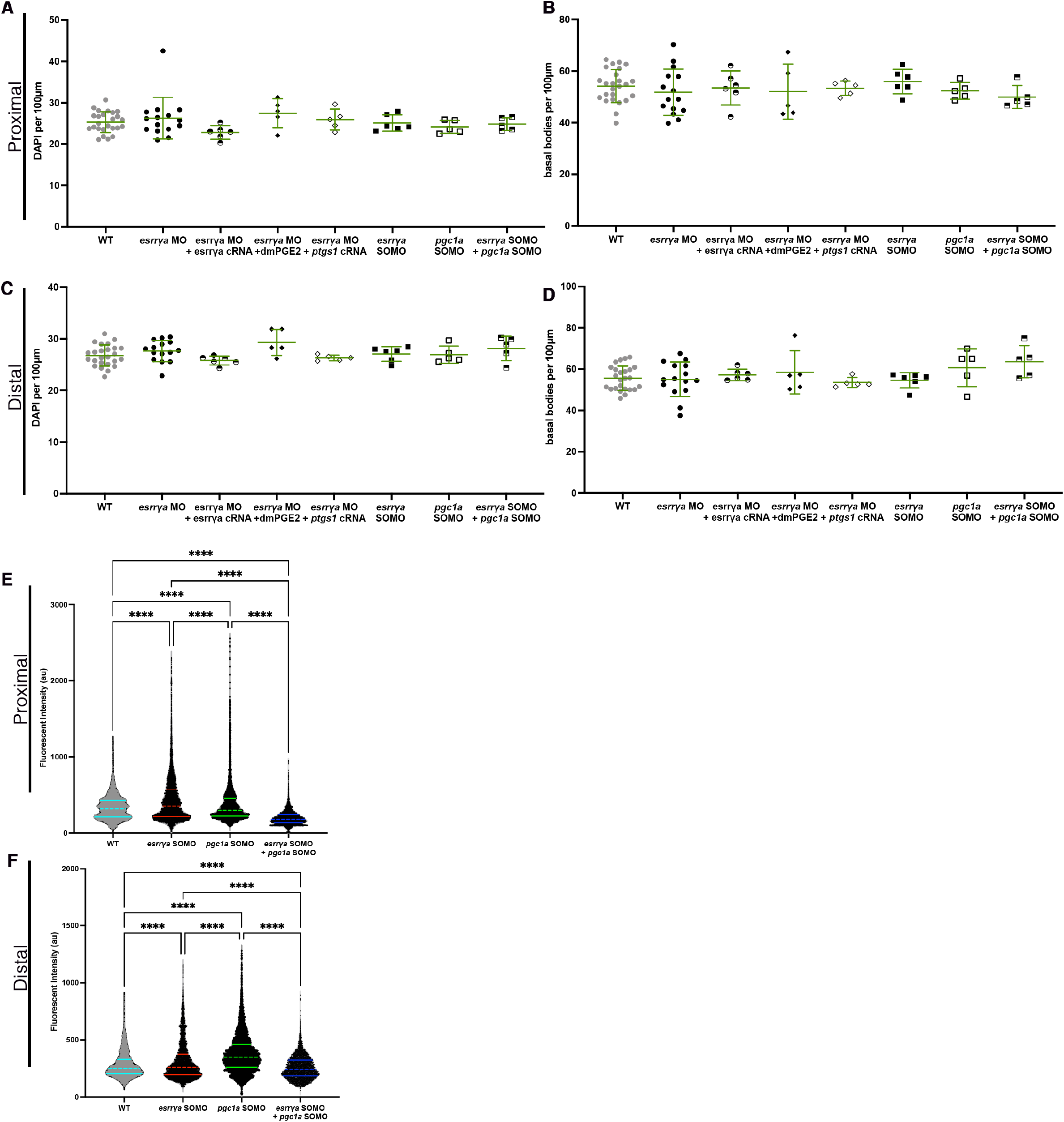
(A) Number of DAPI cells per 100µm in the proximal pronephros. ANOVA statistical test was not significant. (B) Number of basal bodies per 100µm in the proximal pronephros. ANOVA statistical test was not significant. (C) Number of DAPI cells per 100µm in the distal pronephros. ANOVA statistical test was not significant. (D) Number of basal bodies per 100µm in the distal pronephros. ANOVA statistical test was not significant. (E) Fluorescent intensity plots (cilia, alpha tubulin) for the same relative distance in the proximal pronephros at 28 hpf. Representative images seen in Figure 4D. (F) Fluorescent intensity plots (cilia, alpha tubulin) for the same relative distance in the distal pronephros at 28 hpf. Representative images seen in Figure 4D. Data presented on graphs are represented as mean ± SD; * p<0.05, ** p< 0.01 ***p < 0.001 and ****p < 0.0001 (t test or ANOVA).

**Supplemental Figure 5.**
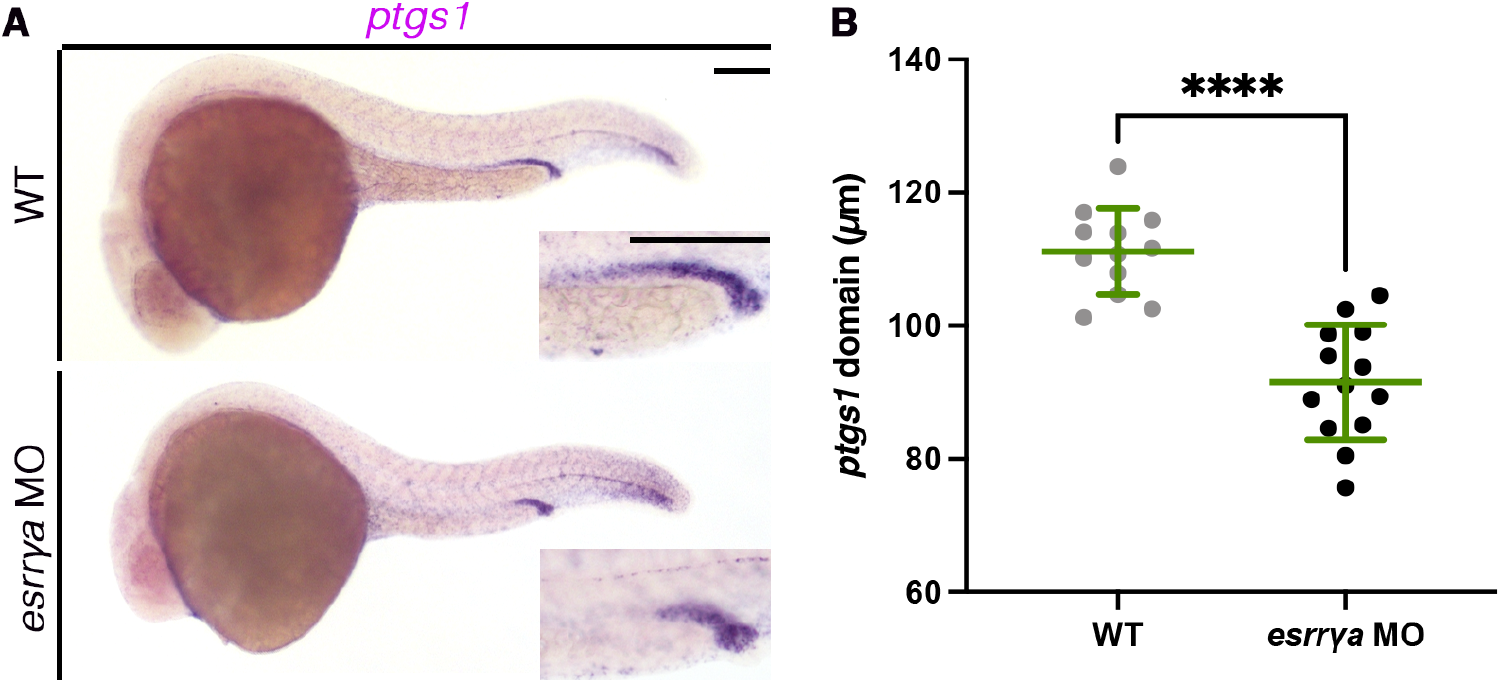
(A) Representative expression of *ptgs1* in WT (top) and *esrrγa* MO (bottom) zebrafish stained via WISH at 24 hpf. Scale bar = 100µm. (B) Length of the *ptgs1* domain in the pronephros at 24 hpf. Data presented on graphs are represented as mean ± SD ****p < 0.0001 (t test or ANOVA).

## Notes

### Competing Interest Statement

The authors have declared no competing interest.

